# Antibody-mediated cell entry of SARS-CoV-2

**DOI:** 10.1101/2023.02.20.529249

**Authors:** Md Golam Kibria, Christy L. Lavine, Weichun Tang, Shaowei Wang, Hailong Gao, Wei Shi, Haisun Zhu, Jewel Voyer, Sophia Rits-Volloch, Keerti, Caihong Bi, Hanqin Peng, Duane R. Wesemann, Jianming Lu, Hang Xie, Michael S. Seaman, Bing Chen

## Abstract

Severe acute respiratory syndrome coronavirus 2 (SARS-CoV-2) enters host cells by first engaging its cellular receptor angiotensin converting enzyme 2 (ACE2) to induce conformational changes in the virus-encoded spike protein and fusion between the viral and target cell membranes. We report here that certain monoclonal neutralizing antibodies against distinct epitopic regions of the receptor-binding domain of the spike can replace ACE2 to serve as a receptor and efficiently support membrane fusion and viral infectivity. These receptor-like antibodies can function in the form of a complex of their soluble immunoglobulin G with Fc-gamma receptor I, a chimera of their antigen-binding fragment with the transmembrane domain of ACE2 or a membrane-bound B cell receptor, indicating that ACE2 and its specific interactions with the spike protein are dispensable for SARS-CoV-2 entry. These results suggest that antibody responses against SARS-CoV-2 may expand the viral tropism to otherwise nonpermissive cell types; they have important implications for viral transmission and pathogenesis.

## Introduction

Enveloped viruses, such as SARS-CoV-2, infect their host cells by first engaging a specific cellular receptor for viral attachment and ultimately by facilitating fusion between the viral and target cell membranes, to deliver the viral genome into the cytoplasm. Membrane fusion is catalyzed by virus-encoded fusion proteins when they refold from a high-energy, metastable prefusion conformational state to a low-energy, stable postfusion state^1–3^. These structural rearrangements are in some cases triggered by binding to the receptor at the cell surface and in others by proton binding at the pH of an endosome, after internalization by endocytosis of the attached viruses^1^. Processing by cleavage and coreceptor interaction may also be involved. The fusion protein of SARS-CoV-2 is its spike (S) protein, which decorates the virion surface^4,5^. The S protein is synthesized as a single polypeptide chain, trimerized and subsequently cleaved by host protease furin into a receptor-binding fragment, S1, and a fusion fragment, S2^6,7^. A receptor-binding domain (RBD) in S1 recognizes the cellular receptor angiotensin converting enzyme 2 (ACE2), and it adopts two different conformations in the S trimer – “up” for a receptor-accessible state and “down” for a receptor-inaccessible state^8,9^. S2 has a transmembrane (TM) segment that anchors the spike in the viral membrane, and another membrane-interacting region, the fusion peptide (FP), which can insert into the target cell membrane^10^. Upon binding of the RBD to ACE2 on a host cell, followed by a second proteolytic cleavage at the S2’ site either by TMPRSS2 (transmembrane serine protease 2) on the cell surface or cathepsin L in endosomes^11,12^, the S trimer undergoes large conformational changes, including dissociation of S1, formation of an extended intermediate that bridges cell and viral membranes, and irreversible refolding of S2 into a postfusion structure^10,13,14^. Formation of the postfusion S2 structure provides the free energy needed to overcome the kinetic barrier for membrane fusion, placing the TM and FP at the same end of the molecule to bring the viral and cellular membranes close together and inducing the two membranes to fuse into a single lipid bilayer^10^.

ACE2 also serves as the cellular receptor for several other coronaviruses, such as SARS-CoV, human coronavirus NL63 and SARS-related bat viruses^11,15–18^. Its interactions with various viral spike proteins have been studied extensively^19–21^. For instance, the binding interface between ACE2 and the SARS-CoV-2 RBD is formed primarily by the N-terminal helix of ACE2 and a gently concave surface of the extended receptor binding motif (RBM) in the RBD, with extensive networks of hydrophilic interactions that account for the affinity and specificity^19,20^. ACE2 binding appears to facilitate dissociation of S1 when a virion binds at the surface of an ACE2 expressing cell, leading to formation of an extended intermediate by S2, which subsequently collapses to induce membrane fusion, when exposed to the mildly acidic pH^22^. Structural studies of the ACE2-bound S trimers have not shown any obvious differences from the structure of the unliganded S trimer in the RBD-up conformation^23,24^, and the mechanism by which ACE2 induces S1 dissociation remains to be determined.

In patients who died with severe COVID-19, SARS-CoV-2 was rapidly disseminated and widely distributed in multiple respiratory and non-respiratory tissues, including those in brain^25^, inconsistent with the expression profile of ACE2 and TMPRSS2^26,27^, raising the possibility of ACE2-independent entry. Indeed, a number of alternative receptors, including CD147, C-type lectins, phosphatidylserine receptors, heparan sulfate and neuropilin-1 (NRP1), have been suggested to facilitate SARS-CoV-2 entry into specific cell types^28^. Other related coronaviruses (CoVs) use different host proteins as their entry receptor: dipeptidyl peptidase 4 (DPP4) by MERS-CoV and related BatCoV-HKU4^29–31^; amino peptidase N (APN) by several alpha-CoVs, such as human coronavirus HCoV-229E^32–34^; the murine carcinoembryonic antigen-related cell adhesion molecule 1 (CEACAM1) by the N-terminal domain of the S protein from murine hepatitis virus (MHV; ref^35,36^). These related spike proteins can be activated by totally different mechanisms that do not require ACE2, although many have rather different receptor-binding domains.

Strong antibody responses against the SARS-CoV-2 spike are induced by natural infection or vaccination and target different epitopic regions of the protein^37–41^. Selected RBD-directed neutralizing antibodies show Fc-gamma receptor (FcγR)-mediated enhancement of virus infection in cell culture, but administration of these antibodies before SARS-CoV-2 infection in animals do not increase infection in vivo^42^. Nevertheless, analysis of COVID-19 patient samples indicates that blood monocytes and lung macrophages, both expressing ACE2 at almost an undetectable-level, are infected by SARS-CoV-2 via an FcγR/antibody-mediated mechanism^43,44^. The infection of these apparently “nonpermissive” cells causes rapid inflammatory cell death that prevents production of infectious viruses but induces systemic inflammation possibly enhancing disease severity in some individuals^44^, suggesting that an ACE2-independent entry pathway may contribute to COVID-19 pathogenesis. We set out to investigate whether or not anti-spike monoclonal antibodies, when present in a membrane-bound form, could replace ACE2 and directly function as an alternative receptor for SARS-CoV-2, aiming to advance our understanding of the viral entry mechanism and pathogenesis.

## Results

### Membrane fusion mediated by soluble IgG in complex with FcγRI

Previous studies have shown that several anti-RBD neutralizing antibodies against SARS-CoV-2 can mediate enhanced infection by pseudotyped viruses of TZM-bl cells expressing either FcγRI or FcγRIIb but lacking ACE2 and TMPRSS2^42^. It is unclear, however, whether the antibody-FcγR complex simply facilitates endocytosis of the attached viruses, which then cross the endosomal membrane by other mechanisms, or the antibody can replace ACE2 and function as an entry receptor directly. We selected eight spike-specific monoclonal antibodies with seven of them isolated from COVID-19 convalescent individuals (C63C8, G32B6, C12A2, S2H97, C63C7, C12C9 and C81D6)^45,46^ and one identified from humanized mice by immunization (SP1-77)^47^. G32B6 and C12A2 target the RBM and directly compete with ACE2 for the RBD binding; C63C8, SP1-77, S2H97 and C63C7 recognize RBD epitopes outside of the ACE2-binding site; C12C9 and C81D6 are NTD-directed antibodies (Fig. 1A). All these antibodies except for C81D6 potently neutralize the original Wuhan-Hu-1 strain. We used a cell-cell fusion assay^14^ to test whether any of these selected antibodies could support membrane fusion in the presence of the high-affinity FcγRI^48^. To create the target cells, we expressed the FcγRI α chain and the common γ chain in HEK293T cells to capture soluble IgG antibodies on the cell surfaces (Fig. 1B; ref^49^). We produced S-expressing cells using the S protein derived from an early variant G614 (lineage B.1)^50^. When the two types of cells were mixed, four anti-RBD antibodies (C63C8, G32B6, C12A2 and S2H97) showed various levels of the fusion activity, which depended on both FcγRI and antibody (Fig. 1C and S1A). The other two RBD-specific antibodies (SP1-77 and C63C7) and the two NTD-directed antibodies (C12C9 and C81D6) showed no detectable activities. Using the same assay, we also detected weak fusion activities in some polyclonal IgG antibodies purified from vaccinated convalescent individuals (Fig. 1D) or serum samples from convalescent individuals recovered during the early stage of the COVID-19 pandemic (Fig. S1B). These results suggest that selected anti-RBD antibodies can support membrane fusion and that such antibodies are present in some infected or vaccinated individuals.

**Figure 1.**
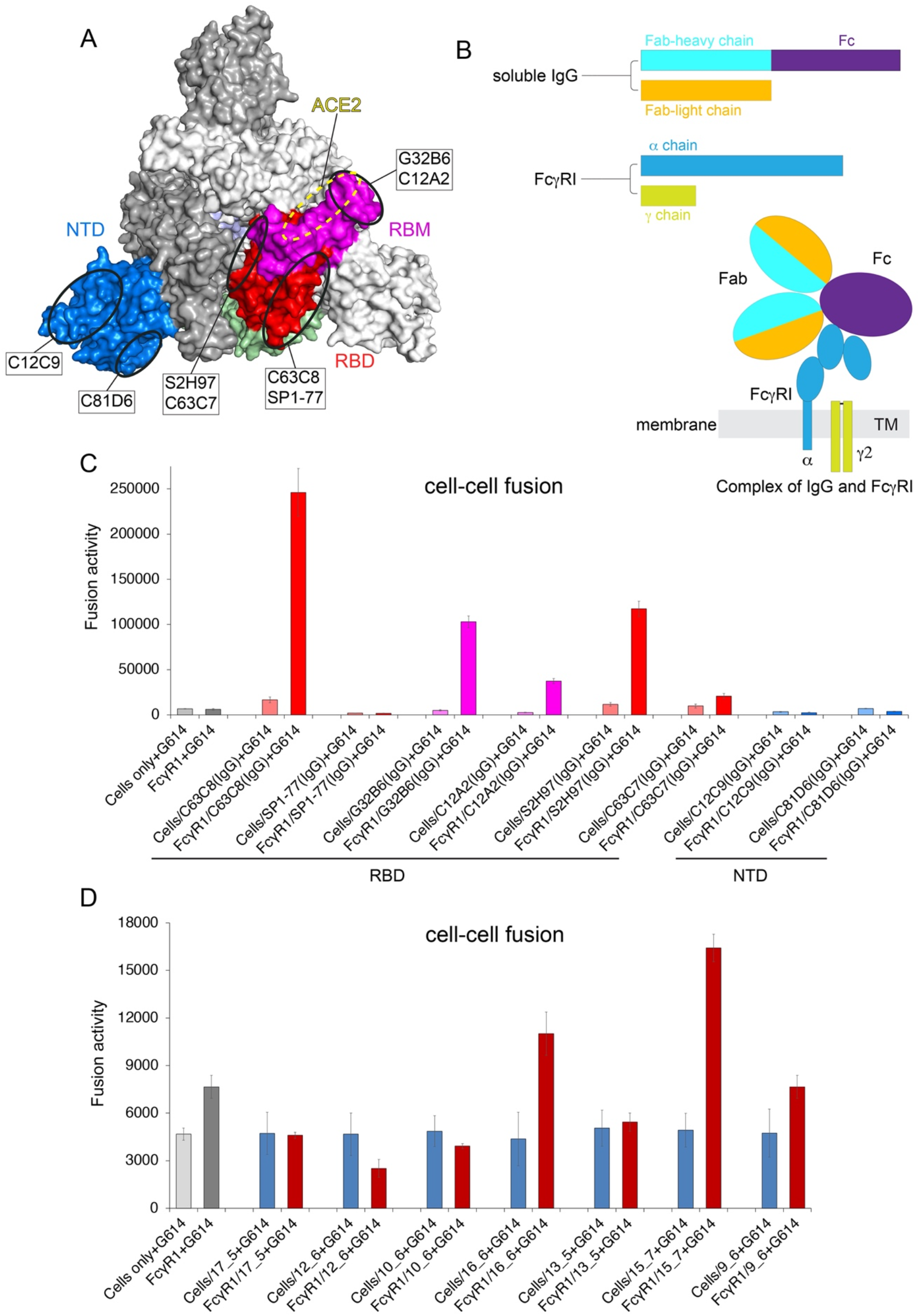
Membrane fusion mediated by spike-specific antibodies. (**A**) Binding site locations of selected monoclonal antibodies and ACE2 binding site. Surface regions of the SARS-CoV-2 spike trimer in a top view targeted by eight selected antibodies on the RBD and NTD are highlighted by ellipses. Various domains of one protomer are colored (RBM in magenta, the rest of RBD in red and NTD in blue); the other two protomers in white and gray, respectively. The ACE2 binding site is marked with a yellow dashed line. (**B**) Schematic representation of antibody IgG (heavy and light chains) and FcγRI (α and γ chains) constructs, as well as a diagram for how FcγRI captures an IgG antibody on the surface of membrane based on the crystal structure PDB ID: 4W4O^74^. (**C**) HEK293T cells with or without expressing FcγRI were decorated with eight selected monoclonal antibodies and tested for membrane fusion with the full-length G614 S protein expressing cells in our standard cell-cell fusion assay. Cell-cell fusion led to reconstitution of α and μ fragments of β-galactosidase yielding an active enzyme and thus the fusion activity was quantified by a chemiluminescent assay. Cells only and cells expressing FcγRI with no antibody added were negative controls. (**D**) The cell-cell fusion assay was used to analyze seven polyclonal IgG antibodies purified from serum samples of vaccinated convalescent individuals reported previously^70^.

### Membrane fusion and viral entry mediated by antibody-ACE2 chimeras

To investigate the mechanism of the antibody-mediated membrane fusion, we next tested whether the catalytic domain of ACE2, which contains the RBD binding site, could be replaced by the antigen-binding (Fab) fragment of a spike-specific IgG antibody for the receptor function. We generated antibody-ACE2 chimeric constructs by fusing the Fab heavy chain with the neck domain, TM anchor and the cytoplasmic tail (CT) of ACE2 (hence mAb-ACE2t), and co-expressed them with their cognate light chain constructs, as depicted in Fig. 2A. We confirmed that expression of these mAb-ACE2t chimeras in HEK293T cells did not lead to increased expression of the endogenous ACE2 (Fig. S2A). We used the cell-cell fusion assay to quantify the fusion activity catalyzed by S proteins derived from both the G614 strain and Omicron subvariant BA.2. When G614 S-expressing cells were mixed with ACE2-expressing cells, they formed large aggregates within 15 minutes, presumably caused by specific interactions between S and ACE2 proteins on the cell surfaces since no obvious association was observed between the cells without expressing these proteins under the same conditions (Fig. S2B). Likewise, SP1-77-ACE2t- and C12C9-ACE2t-expressing cells showed similar aggregates with the G614 S-expressing cells, while C63C8-ACE2t, G32B6-ACE2t, C12A2-ACE2t and S2H97-ACE2t led to smaller aggregates, but C63C7-ACE2t and C81D6-ACE2t gave no obvious aggregates, suggesting most selected antibodies retained their ability to bind the G614 S protein in the form of a chimera with the ACE2 TM region. For the BA.2 S-expressing cells, only ACE2 and SP1-77-ACE2t showed obvious aggregates while the rest of the constructs did not (Fig. S2B).

**Figure 2.**
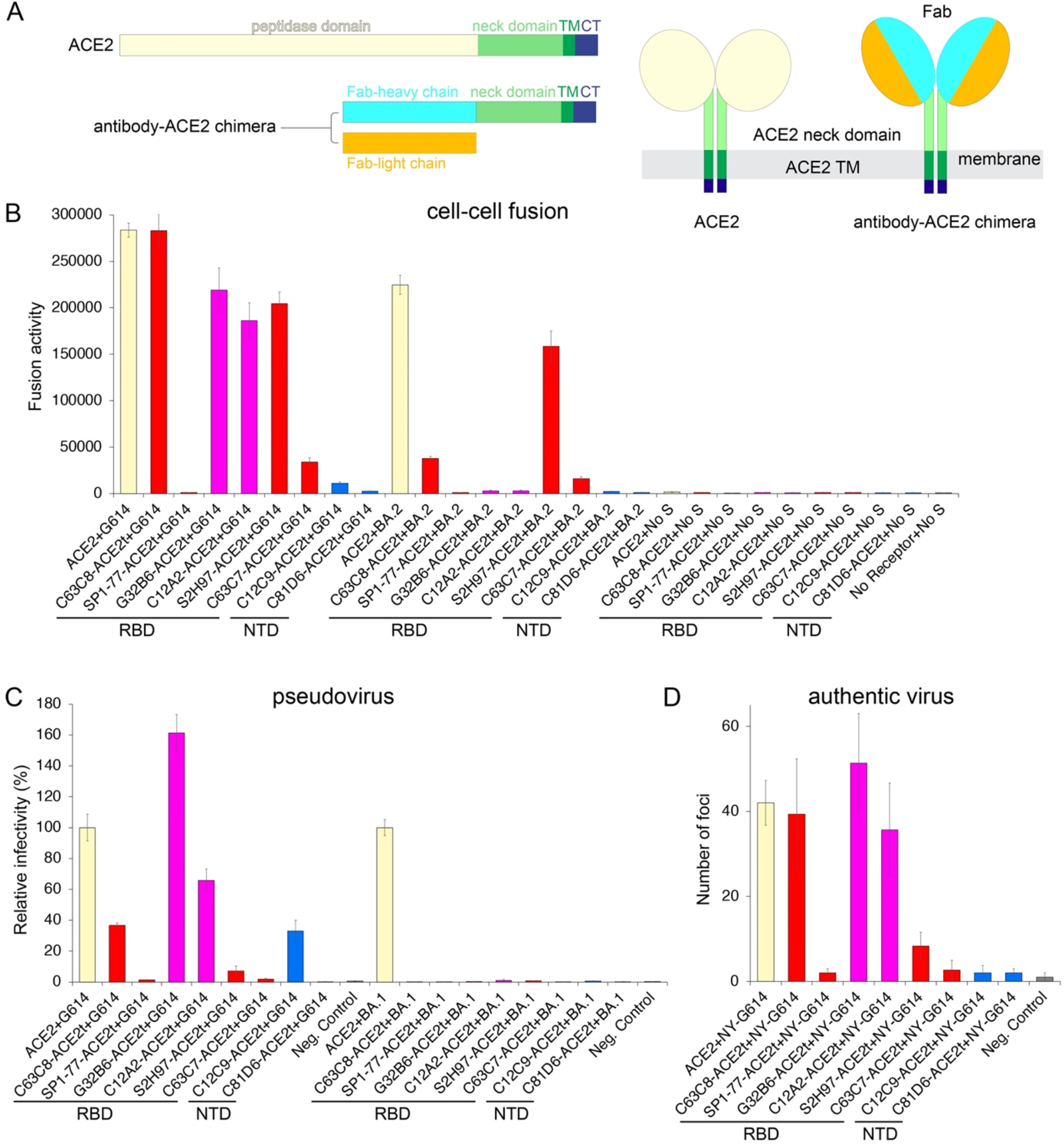
Membrane fusion and viral entry mediated by antibody-ACE2 chimeras. (**A**) Schematic representation of the full-length human ACE2 and design of antibody-based expression constructs. Various segments for ACE2 include: catalytic peptidase domain, neck domain; TM, transmembrane anchor; and CT, cytoplasmic tail. Expression constructs of antibody-ACE2 chimera, the Fab heavy chain of an antibody is fused with the neck domain, TM and CT of ACE2, coexpressed with the Fab light chain. A diagram showing how Fab is presented on the cell surfaces by the ACE2 TM anchor. (**B**) HEK293T cells transfected with eight different antibody-ACE2 chimeric constructs were tested for membrane fusion with the full-length S protein (G614 or Omicron subvariant BA.2) expressing cells in the β-galactosidase-based cell-cell fusion assay. The wildtype ACE2 was a positive control; no receptor/no S a negative control. (**C**) Infection of HEK293T cells transfected with either ACE2 or various antibody-ACE2 chimeric constructs by HIV-based pseudotyped viruses using the full-length G614 and BA.1 S constructs in a single cycle. Empty vector was used as a negative control. (**D**) Virus foci in MDCK cells transfected with either ACE2 or various antibody-ACE2 chimeric constructs followed by infection of the authentic SARS-CoV-2 G614 isolate. Empty vector was used as a negative control.

As expected, ACE2-expressing cells fused very efficiently with the cells expressing either the G614 or Omicron BA.2 S protein (Fig. 2B). Consistent with our observation using IgG antibodies on the FcγRI-expressing cells, five anti-RBD neutralizing antibodies, when anchored in the membrane by the ACE2 TM, also supported membrane fusion with the cells expressing the G614 S. Quantitively, normalized by the amount of plasmid DNA used for transfection, the C63C8-ACE2t chimera was as effective as the wildtype ACE2; G32B6-ACE2t, C12A2-ACE2t and S2H97-ACE2t had 65-77% of the fusion activity by ACE2; C63C7-ACE2t and C12C9-ACE2t showed 12% and 4% activity, respectively; SP1-77-ACE2t and C81D6-ACE2t had no fusion activity (Fig. 2B). C12C9-ACE2t and SP1-77-ACE2t showed the strongest cell-cell association among all eight antibodies, but they had very low or no fusion activity, suggesting that binding to the S protein for attachment alone is not sufficient for membrane fusion. For the cells expressing the BA.2 S protein, the fusion activity of C63C8-ACE2t was only 17% of that by ACE2, consistent with weaker binding of C63C8 to BA.2 S than to G614 S, because of the mutations in this Omicron variant^51^. G32B6-ACE2t and C12A2-ACE2t completely lost their fusion activity since these two antibodies can no longer bind to the BA.2 S trimer^51^. S2H97-ACE2t and C63C7-ACE2t showed 71% and 7% of the ACE2-mediated fusion activity, respectively, not very different from those with the G614 S, because their epitopes in the so-called “cryptic site” on the RBD are relatively conserved and they still retain their binding with the Omicron S proteins^51,52^. SP1-77-ACE2t and C81D6-ACE2t remained inactive in membrane fusion with the BA.2 S-expressing cells, even though SP1-77 potently neutralizes the Omicron variants and C81D6 still binds the purified BA.2 S trimer^47,51^.

To quantify relative expression levels of ACE2 and mAb-ACE2 chimeras, we added a C-terminal GFP tag to all the expression constructs (Fig. S3A). These constructs showed almost the same pattern of antibody dependence in the cell-cell fusion assay as those without the GFP tag (Fig. S3B). When quantified by the GFP expression level in the whole cells, S2H97-ACE2t-GFP, SP1-77-ACE2t-GFP and ACE2-GFP were among the highest expressors and C63C7-ACE2t-GFP and C81D6-ACE2t-GFP the lowest, while others were somewhere in-between (Fig. S3C). When measured by the Fab-expression level on the cell surfaces, S2H97-ACE2t-GFP was the highest and G32B6-ACE2t-GFP the lowest, while others were in-between (Fig. S3D). These results showed that the protein expression levels of the mAb-ACE2 chimeras vary probably due to their intrinsic protein properties, but the expression did not correlate with the membrane fusion activity.

To confirm whether these mAb-ACE2 chimeras can function as an entry receptor for SARS-CoV-2 pseudoviruses and authentic viruses, we first used an HIV-based pseudovirus assay with the full-length G614 S and transiently transfected HEK293T cells. Only C63C8-ACE2t, G32B6-ACE2t and C12A2-ACE2t showed significant activities in supporting infectivity while S2H97-ACE2t was very weak (Fig. 2C). Unexpectedly, C12C9-ACE2t was also positive in this assay. None of the antibodies showed fusion activity with the Omicron subvariant BA.1 S. Next, using nonpermissive MDCK cells transiently transfected with these mAb-ACE2 chimeric constructs, infection of the authentic SARS-CoV-2 G614 virus resulted in formation of positive virus foci with C63C8-ACE2t, G32B6-ACE2t, C12A2-ACE2t and S2H97-ACE2t, while the rest four constructs, including C12C9-ACE2t, showed no or only the background signals (Fig. 2D and S4). The discrepancies among the cell-cell fusion assay and the two types of viral infectivity assays may be due to different surface-expression levels of the receptor-like constructs and lack of other SARS-CoV-2 components with cell-cell fusion and pseudovirus infection. Nonetheless, we have clearly demonstrated that the Fab fragments of several RBD-specific antibodies can replace the catalytic domain of ACE2 to function as an entry receptor for SARS-CoV-2.

### Membrane fusion and viral entry mediated by B cell receptors

Since the antibody-FcγRI complex can support membrane fusion, we predicted that the C-terminal portion of ACE2 in the chimeric constructs would not be required for the receptor function either. We therefore reconstructed these antibodies in a membrane-bound B cell receptor (BCR) form by fusing the Fab heavy chain with a segment contain the Fc region, TM and CT of a BCR (Fig. 3A). We also confirmed that the expression levels of these BCR constructs in HEK293T cells also varied, but their production did not increase the expression of the endogenous ACE2 (Fig. S5). As predicted, C63C8/BCR, G32B6/BCR, C12A2/BCR and S2H97/BCR indeed supported membrane fusion with the cells expressing the G614 S and S2H97/BCR was also active with the BA.2 S protein (Fig. 3B). In the viral infectivity assays, only C63C8/BCR was consistently positive with both the pseudoviruses and authentic viruses with the G614 S (Fig. 3C, 3D and S6). Thus, a selected B cell receptor, totally unrelated to ACE2, can also serve as an entry receptor for SARS-CoV-2, although how the Fab of a receptor-like antibody is presented on a cell surface clearly has an important impact on viral entry.

**Figure 3.**
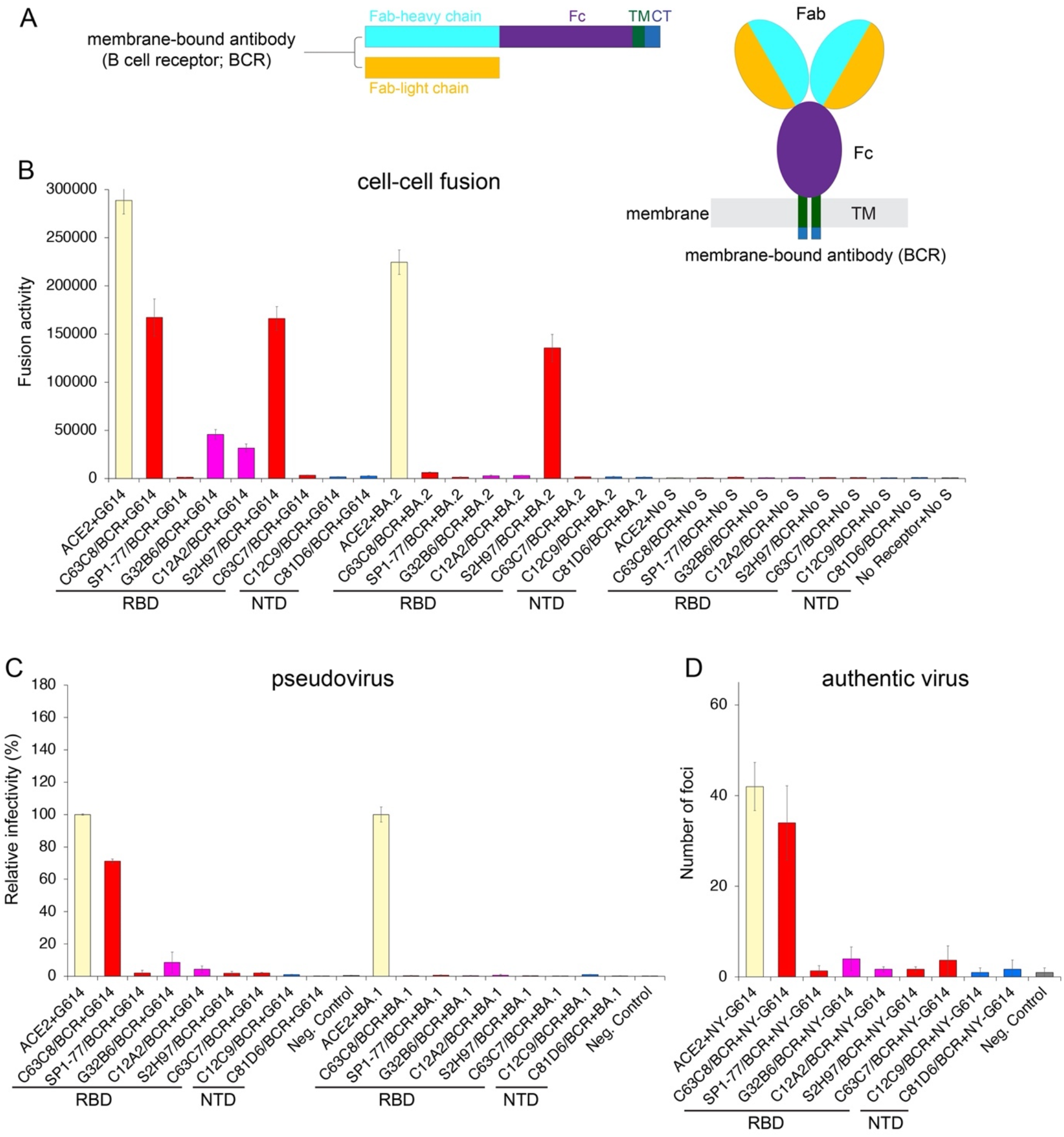
Membrane fusion and viral entry mediated by BCRs. (**A**) Schematic representation of the membrane-bound B cell receptor (BCR) expression constructs and a diagram showing how Fab is presented on the cell surfaces by the BCR TM anchor. (**B**) HEK293T cells transfected with eight different membrane-bound BCR constructs were tested for membrane fusion with the full-length S protein (G614 or Omicron subvariant BA.2) expressing cells in the β-galactosidase-based cell-cell fusion assay. The wildtype ACE2 was a positive control and no receptor/no S a negative control. (**C**) Infection of HEK293T cells transfected with either ACE2 or various BCR constructs by HIV-based pseudotyped viruses using the full-length G614 and Omicron BA.1 S constructs in a single cycle. Empty vector was used as a negative control. (**D**) Virus foci in MDCK cells transfected with either ACE2 or various antibody-ACE2 chimeric constructs followed by infection of the authentic SARS-CoV-2 G614 isolate. Empty vector was used as a negative control.

### Inhibition of SARS-CoV-2 entry mediated by receptor-like antibodies

We analyzed inhibition by protease inhibitors and neutralizing antibodies of SARS-CoV-2 entry using the pseudovirus assays. The S protein is cleaved at the S2’ site by TMPRSS2 for cell surface entry, or by cathepsin L after endocytosis for endosomal entry^11,12,53^. For the ACE2-mediated entry into HEK293 cells, it was only partially sensitive to the cathepsin L inhibitor E-64d when TMPRSS2 was not overexpressed (Fig. 4A, 4B and S7). The infectivity of the HIV-based pseudoviruses using the full-length G614 S increased substantially when TMPRSS2 was overexpressed and it became partially sensitive to the TMPRSS2 inhibitor Camostat, but no longer to E-64d anymore, consistent with a previous report that the TMPRSS2 pathway is preferred when available^54^. Inhibition of C63C8-ACE2t and C63C8/BCR mediated entry by the two inhibitors showed a very similar pattern (Fig. 4A and S7), suggesting that the entry pathways were not altered when C63C8 served as an entry receptor. In contrast, the G32B6-ACEt mediated entry was not sensitive to either inhibitor, but G32B6/BCR showed a profile similar to that by ACE2 (Fig. 4B and S7). These results suggest that the antibody-based receptors largely preserve the two entry pathways and show similar sensitivities to the host protease inhibitors, but the configuration in which a receptor-like antibody is presented on the cell surfaces may change the sensitivity to the host proteases.

**Figure 4.**
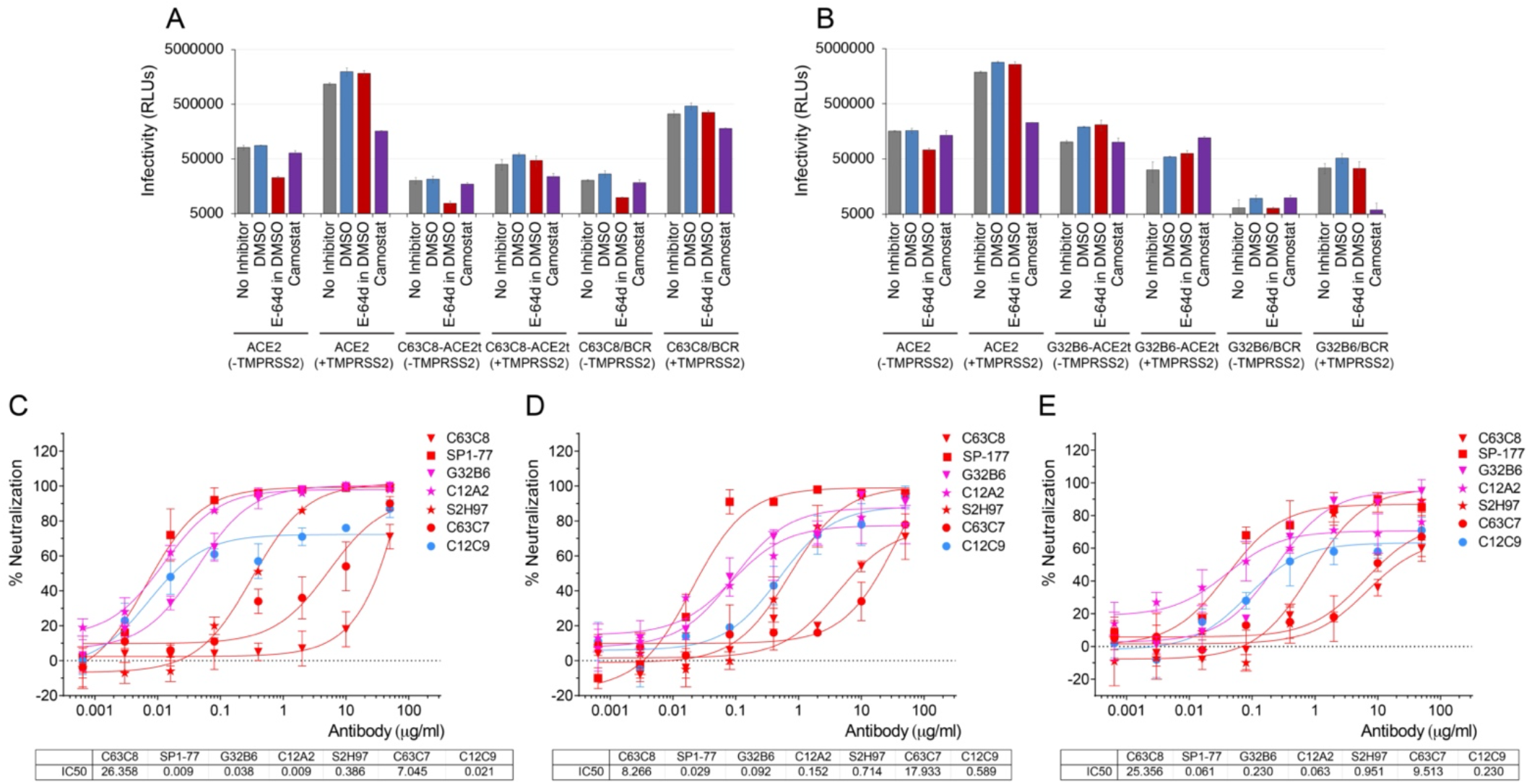
Inhibition of SARS-CoV-2 entry mediated by receptor-like antibody. (**A**) and (**B**) Inhibition of viral infectivity by protease inhibitors. Pseudovirus (G614 S) infection of HEK293T cells transfected with ACE2 or antibody constructs with/without TMPRSS2 were treated with either E-64d (a cathepsin L inhibitor) or Camostat (TMPRSS2 inhibitor). DMSO, organic solvent used to dissolve E-64d. (**C**-**E**) Antibody neutralization of pseudoviruses containing the G614 S protein was determined using IgG antibodies, C63C8, SP1-77, S2H97 and C63C7 in red; G32B6 and C12A2 in magenta; and C12C9 in blue.

To minimize the variations among different transient transfections, we generated stably transfected cell lines expressing C63C8-ACE2t and C63C8/BCR, and used them as the target cells with the HEK293/ACE2 cell line as a reference to analyze antibody neutralization. We first confirmed that both cell lines are susceptible to infection by several early SARS-CoV-2 variants of concern, but not by the recent Omicron subvariant in a different, MLV-based pseudovirus assay, as expected (Fig. S8). The HIV-based pseudoviruses were less sensitive to most antibodies tested when using the antibody receptor for entry than using ACE2 (Fig. 4C and 4D). Surprisingly, G32B6 and C12A2, which target the ACE2 binding site and are believed to neutralize by competing with ACE2 for the RBD binding, still potently neutralized the virus even when C63C8, recognizing a noncompeting epitope, served as the receptor, suggesting that the neutralization mechanism by these antibodies is more complicated than simply blocking the receptor binding.

We have previously shown that soluble ACE2 can trigger the conformation changes of the purified full-length S trimer and induce S1 dissociation^10^. To assess whether the ability to induce S1 shedding correlates with the receptor function of an antibody, we incubated all eight selected IgG antibodies with the purified S trimer, and analyzed by gel filtration chromatography and SDS-PAGE. As shown in Fig. S9, the prefusion G614S trimer purified in detergent was very stable in solution, but it could be converted into the postfusion conformation when treated with soluble ACE2. C63C8 and S2H97, but not G32B6 and C12A2, induced conformational changes of the S trimer, although they can all support membrane fusion. This observation was further confirmed with the purified S trimer reconstituted in lipid bilayer using nanodiscs (Fig. S9). Thus, the ability of an antibody to induce S1 dissociation using purified proteins does not predict its ability to serve as an entry receptor for SARS-CoV-2.

## Discussion

Our finding that selected monoclonal neutralizing antibodies that target the RBD of SARS-CoV-2 spike can replace ACE2 and function as an entry receptor provides a plausible explanation for how the virus infects those apparently nonpermissive cells (e.g., blood monocytes and tissue-resident macrophages) in COVID-19 patients^43,44^. Although previous studies show that monocyte infection relies on FcγRs to uptake antibody-decorated viruses by endocytosis^44^, how the virus crosses the endosomal membrane and enters the cytosol of these cells remains unclear. Our data demonstrate that certain RBD-directed IgG antibodies, when captured by FcγRI on the surfaces of target cells, can serve as an entry receptor, independent of ACE2, and efficiently support S-catalyzed membrane fusion. The new information fills an important gap in our understanding of the COVID-19 pathogenesis. Based on the very small number (total 8) of monoclonal antibodies that we have tested, 4 out of 6 RBD-directed antibodies, but not the NTD-specific antibodies, can function efficiently as an entry receptor, even for the authentic virus (summarized in Table S1). Moreover, such a membrane-fusion activity can also be detected in purified polyclonal IgG antibodies or serum samples from some convalescent individuals (Figs. 1D and S1B). The clinical significance of the presence of these receptor-like antibodies needs to be determined in future studies with a much larger sample size than what presented here.

As for the molecular mechanism of viral entry, it is surprising that the specific binding of the RBD to ACE2 is not required for membrane fusion catalyzed by the SARS-CoV-2 spike, although binding to a specific entry receptor is necessary to activate the pH-independent class I viral fusion proteins that have been characterized^1^. The receptor-like antibodies identified in this study target three distinct regions of the RBD, including the exposed surface of the domain in the RBD-down conformation (C63C8), the ACE2-binding site (G32B6 and C12A2) and the cryptic site fully exposed only in the RBD-up conformation (S2H97), suggesting that specific interactions between the RBD and ACE2 are not essential for triggering the S protein to undergo conformational changes driving the membrane fusion, nor are any portions of ACE2. The underlying mechanism of how different types of antibodies, while all neutralizing, can activate the S protein and achieve the entry receptor’s function is currently unknown. It is clear that the nonneutralizing antibody C81D6 cannot function as a receptor probably because it does not bind to the membrane-bound S trimer to allow initial attachment. However, attachment alone is not sufficient since both C12C9 and SP1-77 support strong cell-cell attachment, which does not correlate with their low or no fusion activities. Although SP1-77 consistently shows no receptor-like function under all conditions, probably because it neutralizes by locking the S trimer in the prefusion conformation and preventing S1 dissociation^47^, the receptor function does not appear to correlate with the ability to induce S1 dissociation either, at least with purified proteins. In addition, neutralization is not a major determinant either because the strongest receptor-like antibody C68C3 is weakly neutralizing. Furthermore, the observation that the receptor-like antibodies can be anchored to the membrane in three distinct formats suggests that the precise orientation and distance to the membrane of the RBD-binding surface of a receptor are not crucial for membrane fusion. Thus, the SARS-CoV-2 spike appears to have evolved to use diverse strategies to activate its fusogenic potential, suggesting that the virus may be able to jump to other species more easily and more rapidly than previously recognized, as apparently there is no need to cultivate a specific “receptor-triggering” mechanism through a long period of coevolution with a new host. It remains to be seen whether or not this notion is applicable to other coronaviruses or enveloped viruses.

Since the beginning of the COVID-19 pandemic, concerns have been raised that antibodies induced by natural infection or vaccination may cause antibody-dependent enhancement (ADE) of infectivity and virulence^55,56^. ADE is well-established for dengue virus infection, initially based on the epidemiological evidence and subsequently explained by Fc-gamma receptor (FcγR)-mediated infection of leukocytes^57–60^. Our results suggest that at least two types of cells, previously thought as nonpermissive, can be infected by SARS-CoV-2 and they are FcγR1-expressing cells, such as blood monocytes and lung macrophages, and B cells that express receptor-like BCRs. Conceivably, the existing receptor-like antibodies could lead to SARS-CoV-2 infection of other “nonpermissive” cells expressing no or low levels of ACE2, thereby facilitating rapid spread to other non-respiratory tissues^25^. It may also help explain the recent report that reinfection by SARS-CoV-2 increases risks of all-cause mortality and adverse health outcomes^61^. Nevertheless, the real-life scenario is likely very complicated since blood monocytes and lung macrophages have already been shown not to support productive infection, and other cell types may not support active viral replication either for other reasons, such as host restriction factors^62^.

Finally, all receptor-like antibodies identified in this study are neutralizing. They potently block viral infection when in the soluble form, but can also potentially promote infection of those otherwise nonpermissive cells, when in a membrane-bound form. Emerging SARS-CoV-2 variants, such as the latest Omicron subvariants, cause reinfection in convalescent individuals or breakthrough infection in the vaccinated population primarily by evading pre-existing neutralizing antibodies^63,64^, which are not all receptor-like and which wane within several months^65,66^. The new viruses would also lose the ability to gain entry using the pre-existing receptor-like antibodies, which would not bind the mutated spikes. Thus, the receptor-like antibodies may contribute little to transmission of waves of new SARS-CoV-2 variants, consistent with lack of population-wide epidemiological evidence for ADE to the transmissibility in the vaccinated population.

## Materials and Methods

### Expression constructs of antibody-ACE2 chimeras (mAb-ACE2ts) and BCRs (mAb/BCRs)

A codon-optimized intact human ACE2 (aa 1–805) gene was synthesized and cloned into pCMV-IRES-puro (Codex BioSolutions, Inc, Gaithersburg, MD), as described^23^. Expression constructs for selected monoclonal antibodies C63C8, G32B6, C12A2, C63C7, C12C9, C81D6 and SP1-77 were previously described^45,47^. A synthetic gene for S2H97 was generated by Twist Bioscience (South San Francisco, CA) based on the published sequences^46^. The expression constructs for antibody SP1-77 were kindly provided by Drs. Sai Luo and Frederick Alt (Boston Children’s Hospital). Antibody-ACE2 chimeric (mAb-ACE2t) constructs were created by replacing the catalytic peptidase domain of ACE2 (aa 1-599) in the full-length ACE2 construct with the Fab heavy chain fragment (VH and CH1) of various antibodies, using the standard PCR procedure and ClonExpress II One Step cloning kit (C112-02, Vazyme, China). Membrane-bound B cell receptor (BCR) constructs were produced by fusing the transmembrane (TM) and cytoplasmic tail (CT) segment of human IgG1 (QLEESCAEAQDGELDGLWTTITIFITLFLLSVCYSATVTFFKVKWIFSSVVDLKQTIIPDYRNMIGQGA) to the C-terminal end of the IgG antibodies. All constructs were confirmed by DNA sequencing.

### Cells and virus

HEK293T-derived stable cell lines expressing ACE2, C63C8-ACE2t and C63C8/BCR were created following our published protocol^67^. Briefly, 8×10^5^ HEK293T cells in 2 ml of DMEM containing 10% FBS and no antibiotics were seeded on a 6 well-plate and incubated for overnight. The cells were transfected with the expression constructs using Lipofectamine 3000 (Life Technologies, Grand Island, NY) following a protocol recommended by manufacturer. 24 hours post-transfection, the transfected cells were transferred into a medium containing DMEM, 10% FBS and 1 μg/ml puromycin for selection. Single colonies were picked in 2-3 weeks, and transferred into 24-well plates in the same selective medium. Protein expression was confirmed by western blot. Positive clones were expanded, frozen and stored in liquid nitrogen.

Madin-Darby Cannie Kidney (MDCK) cells, nonpermissive to SARS-CoV-2, were used to test the receptor-like antibody constructs with the authentic virus. MDCK cells obtained from ECACC (#84121903) were maintained in Gibco™ high-glucose Dulbecco’s modified Eagle’s medium (DMEM) supplemented with 5% fetal bovine serum (FBS), 1% GlutaMAX™, 1% penicillin/streptomycin, 10 mM HEPES pH 7.3. TMPRSS2-E6 cells obtained from BPS Bioscience (#78081) were maintained in the same DMEM growth medium supplemented with 10% FBS and additional 3 μg/ml of Puromycin and 1% Na pyruvate. The SARS-CoV-2 G614 seed virus (the clinical isolate New York-PV09158/2020; ATCC #NR-53516) was obtained through BEI Resources (Manassas, VA) and amplified in TMPRSS2-E6 cells. Aliquoted virus was stored in a secured −80 °C freezer until use. Virus was titrated in TMPRSS2-E6 using an ELISA-based 50% tissue culture infectious dose (TCID_50_) method^68,69^. All experiments involving infectious SARS-CoV-2 were performed in an FDA Animal Biosafety Level-3 (ABSL-3) laboratory equipped with advanced access control devices and by trained personnel equipped with powered air-purifying respirators.

### Purification of monoclonal and polyclonal IgG antibodies

To produce monoclonal IgG antibodies, Expi293F cells (ThermoFisher Scientific, Waltham, MA) at a density of 2.5×10^6^ cells/ml were transiently transfected with the IgG heavy chain expression construct and its cognate light chain construct at a concentration of 0.45 μg/ml and 0.54 μg/ml culture, respectively, using Polyethylenimine (PEI). Five days posttransfection, the cell supernatant was collected and soluble IgG was purified using affinity chromatography (GammaBind Plus Sepharose, GE HealthCare, Chicago, IL). Purified IgG was buffer-exchanged in PBS, concentrated and stored at −80 °C. Polyclonal IgG antibodies, from serum samples of vaccinated convalescent individuals reported previously^70^, were purified using Pierce Protein A Agarose (ThermoFisher Scientific).

### Cell-cell fusion assay

The cell-cell fusion assay, based on the α-complementation of E. coli β-galactosidase, was carried out to quantify the fusion activity mediated by various receptor-like antibody constructs, as described^14^. Briefly, the full-length SARS-CoV 2 (G614 or Omicron BA.2) spike construct (10 μg) and the α fragment of E. coli β-galactosidase construct (10 μg) were transfected in HEK293T cells using PEI (80 μg) in 100 mm Petri dishes to prepare S-expressing cells. HEK293T cells were transfected with the full-length human ACE2 construct (5 μg) and empty vector (5 μg), or Fab heavy chain-ACE2t construct (5 μg) and antibody light chain (5 μg), or BCR heavy chain (5 μg) and antibody light chain (5 μg), together with the ω fragment of E. coli β-galactosidase construct (10 μg), to produce receptor-expressing cells. After a 24-hour incubation at 37 °C, the cells were detached using PBS and resuspended in complete DMEM medium. 50 μl S-expressing cells (1.0×10^6^ cells/ml) were mixed with 50 μl receptor-expressing cells (1.0×10^6^ cells/ml) to allow the cell-cell fusion to proceed at 37 °C for 4 hours. Cell-cell fusion activity was quantified using a chemiluminescent assay system, Gal-Screen (Applied Biosystems, Foster City, CA), following the standard protocol recommended by the manufacturer. The substrate was added to the mixture of the cells and allowed to react for 90 minutes in dark at room temperature. The luminescence signal was recorded with a Synergy Neo plate reader (Biotek, Winooski, VT).

To prepare FcγR1 expressing cells, the full-length FcGR1A (SC119841, OriGene Technologies, Inc, MD) and FcER1G (SC117594, OriGene Technologies, Inc, MD) constructs (5 μg of each), together with the ω fragment of E. coli β-galactosidase construct (10 μg), were transfected in HEK293T cells. For cell-cell fusion, 50 μl FcγR1-expressing cells (1.0×10^6^ cells/ml) were first incubated with various IgG antibodies (25 μg/ml) at 37 °C for 30 min and subsequently mixed with 50 μl of S-expressing cells to allow cell-cell fusion to proceed at 37 °C for 4 hours. Cell-cell fusion activity was quantified as described above.

### Western Blot

Western blot was performed using an anti-ACE2 antibody (MAB10823, R&D system, MN) following a protocol described previously^14^. Briefly, samples were prepared from cell pellets expressing mAb-ACE2t or mAb/BCR constructs. Cell lysates were resolved in 4-15% Mini-Protean TGX gel (Bio-Rad, Hercules, CA) and transferred onto PVDF membranes. Membranes were blocked with 3% skimmed milk in PBST overnight at 4 °C and incubated with an anti-ACE2 antibody at a final concentration of 1μg/ml for another 90 min at room temperature. Alkaline phosphatase conjugated anti-Rabbit IgG (1:5000) (Sigma-Aldrich, St. Louis, MO) was used as a secondary antibody. Proteins were visualized using one-step NBT/BCIP substrates (Promega, Madison, WI). HRP conjugated anti-beta antibody (sc-47778, Santa Cruz Biotechnology Inc., Dallas, TX) was used to detect expression of β-actin as the sample loading control.

### Flow cytometry

HEK293T cells transfected by ACE2, mAb-ACE2t, mAb-ACE2t-GFP and mAb/BCR and used for the cell-cell fusion assays were quantified by flow cytometry for the surface presentation of various Fab fragments. After washing by PBS, 1×10^6^ cells in 100 μl were stained with APC conjugated anti-human F(ab’)2 fragment specific antibody (Jackson ImmunoResearch, West Grove, PA) at a dilution of 1:100 and incubated for 1 hour at room temperature. After three additional washes with 1% BSA in PBS, the cells were resuspended in 30 μl of 1% BSA in PBS. Cells were run through an Intellicyt iQue Screener Plus flow cytometer. A blue laser with a wavelength of 488 nm was used to concurrently detect the degree of GFP expression. The flow cytometry assays were repeated two times with essentially identical results.

### HIV-based pseudovirus assay

Neutralizing activity against SARS-CoV-2 pseudovirus was measured using a single-round infection assay in HEK293T target cells expressing either wildtype ACE2 or transiently transfected to express antibody-ACE2 chimeras or BCRs. Pseudotyped virus particles were produced in 293T/17 cells (ATCC) by co-transfection of plasmids encoding codon-optimized SARS-CoV-2 full-length S construct (D614G) or Omicron BA.1, packaging plasmid pCMV ΔR8.2, and luciferase reporter plasmid pHR’ CMV-Luc. Spike, packaging and luciferase plasmids were kindly provided by Drs. Barney Graham and Tongqing Zhou (Vaccine Research Center, NIH). For neutralization assays, serial dilutions of monoclonal antibodies (mAbs) were performed in duplicate followed by addition of pseudovirus. Pooled serum samples from convalescent COVID-19 patients or pre-pandemic normal healthy serum (NHS) were used as positive and negative controls, respectively. Plates were incubated for 1 hour at 37°C followed by addition of 293/ACE2 or 293T cells transiently transfected with antibody-ACE2 chimeras or BCRs as target cells (1×10^4^/well). Wells containing cells + pseudovirus (without sample) or cells alone acted as positive and negative infection controls, respectively. Assays were harvested on day 3 using Promega BrightGlo luciferase reagent and luminescence detected with a Promega GloMax luminometer. Titers are reported as the concentration of mAb that inhibited 50% or 80% virus infection (IC_50_ and IC_80_ titers, respectively). All neutralization experiments were repeated twice with similar results.

Inhibition of viral infectivity by protease inhibitors was measured by pre-incubating target cells (1×10^4^ cells/well) with predetermined optimal concentrations of the protease inhibitors E-64d (25μM in DMSO), or Camostat (100μM) for 2 hours. Cells were also incubated with either DMSO (25μM, as a negative control for E-64d) or culture medium (cells only). Following incubation, the target cells were plated in Poly-L-Lysine coated plates containing serially diluted G614 or BA.1 pseudovirus. Wells containing cells + pseudovirus (no protease inhibitors or DMSO) or cells alone acted as positive and negative infections controls, respectively. Assays were harvested on day 3 using Promega BrightGlo luciferase reagent and luminescence was detected with a Promega GloMax Navigator luminometer. Results are reported as Relative Luciferase Units (RLUs). All virus infectivity experiments were repeated twice with similar results.

### MLV-based pseudovirus assay

Murine Leukemia Virus (MLV) particles (plasmids of the MLV components kindly provided by Dr. Gary Whittaker at Cornell University and Drs. Catherine Chen and Wei Zheng at National Center for Advancing Translational Sciences, National Institutes of Health), pseudotyped with various SARS-CoV-2 S protein constructs, were generated in HEK293T cells, following a protocol described previously for SARS-CoV^71,72^. To enhance incorporation of S protein into the particles, the C-terminal 19 residues in the cytoplasmic tail of each S protein were deleted. To prepare for infection, 7.5×10^3^ of HEK 293 cells, stably transfected with a full-length human ACE2 expression construct, in 15 μl culture medium were plated into a 384-well white-clear plate coated with poly-D-Lysine to enhance the cell attachment. On day 2, 12.5 μl of MLV pseudoviruses for each variant were added into each well pre-seeded with HEK293-ACE2 cells. The plate was centrifuged at 114 xg for 5 min at 12°C. After incubation of the pseudoviruses with the cells for 48 hr. Luciferase activities were measured with Firefly Luciferase Assay Kit (CB-80552-010, Codex BioSolutions Inc).

### ACE2 Transfection and cell infection by authentic virus

For mAb-ACE2t or mAb/BCR expression, the heavy chain and light chain constructs were pre-mixed at 1:1 (w/w). The positive control contained the plasmid expressing wild type ACE2 pre-mixed with the empty vector at 1:1 (w/w). The negative control contained just the empty vector. A total of 13.6 μg of empty vector or mixed constructs (6.8 μg/each plasmid) was then used to prepare the transfection mixture with jetPRIME transfection reagent (Polyplus #101000046) for transfection. The transfection mixtures were added to MDCK pre-seeded in 24-well tissue culture plates. Six hours later, transfection mixtures were removed and MDCK cells were then incubated in fresh growth medium at 37°C, 5% CO_2_ for another 2 days. After aspiration of the growth media, transfected MDCK cells were infected with live SARS-CoV2 G614 at 2×10^5^ TCID_50_/350 μl/well at 37 °C, 5% CO_2_ for 3 hours. The inoculum was removed, and cells were overlaid with 1.2% of Avicel (FMC BioPolymer #CL-611) in 1X EMEM (Quality Biological #115-073-101) containing 2% FBS and 1% penicillin/streptomycin. After continuous incubation at 37°C, 5% CO_2_ for 2 days, cells were fixed with 10% neutral buffered formalin at room temperature for 20 min followed by permeabilization with 0.1% Triton X-100 (sigma #9036-19-5) for another 20 min. After washing, infected cells were probed with anti-nucleocapsid rabbit monoclonal antibody (SinoBiol #40143-R001, 1:6000) at 4°C overnight followed by peroxidase-conjugated goat anti-rabbit secondary antibody (SeraCare #5220-0336, 1:2000) at room temperature for another 1 hour. Foci were developed using KPL TrueBlue substrate (SeraCare #5510-0030) and images were acquired using AID vSpot ELISPOT reader (Autoimmun Diagnostika GmbH, Strassberg, Germany).

### S1 shedding induced by ACE2 or antibodies

The purified full-length G614 spike protein was mixed with soluble ACE2 protein or IgG antibodies with a molar ratio of 1:6 in buffer A containing 25 mM Tris-HCl, pH 7.5, 150 mM NaCl, 0.02% DDM, and incubated at room temperature for 30 min. The mixture was resolved by gel filtration chromatography on a Superose 6 10/300 column (GE Healthcare) in buffer A. For S1 shedding from the S trimer in membrane, the S protein was first reconstituted in lipid nanodiscs as described previously^10^. Briefly, the spike protein and the soy extract polar lipid (Avanti, Birmingham, AL) were mixed and incubated on ice for 30 min. The csMSP2N2 was added to the mixture with a spike:csMSP2N2:lipid molar ratio of 1:8:700 and incubated on ice for another 30 min. Bio-beads SM2 (Bio-Rad, Hercules, CA) were added to remove detergents from the mixture and initiate the reconstitution with rotation at 4 °C overnight. The S-nanodisc sample was incubated with antibody C63C8 or G32B6 with a molar ratio of 1:6 in buffer B containing 25 mM Tris-HCl, pH 7.5, 150 mM NaCl, and incubated at room temperature for 30 min. The mixture was resolved by gel filtration chromatography on a Superose 6 10/300 column in buffer B.

### Negative stain EM

To prepare grids, 4 μl of freshly purified peak fraction sample was adsorbed to a glow-discharged carbon-coated copper grid (Electron Microscopy Sciences, Hatfield, PA), washed with deionized water and stained with freshly prepared 1.5% uranyl formate. Images were recorded at room temperature on the Phillips CM10 transmission electron microscope with a nominal magnification of 52,000×. Particles were auto-picked and 2D class averages were generated using RELION 4.0.0^73^.

## Acknowledgments

We thank S. Harrison for critical reading of the manuscript, S. Luo and F.W. Alt for providing the expression constructs of antibody SP1-77. This work was supported by COVID-19 Awards by MassCPR (to B.C., D.R.W., and M.S.S.), Fast grants by Emergent Ventures (to B.C. and D.R.W.), and NIH grants AI147884 (to B.C.), AI141002 (to B.C.), AI127193 (to B.C. and James Chou) and AI39538, AI170715, AI165072, and AI169619 (to D.R.W), and a CBER/FDA intramural grant (to H.X.)

## Author Contribution

B.C., M.G.K. and H.G. conceived the project. M.G.K. designed and constructed all receptor-like antibody expression plasmids, purified antibodies and performed cell-cell fusion assays with help from J.V., H.P. and S.R.V.. C.L.L. and M.S.S performed the infectivity, protease inhibition and antibody neutralization assays using the HIV-based pseudoviruses. W.T. and H.X. carried out the infectivity assay using the authentic SARS-CoV-2. J.L. and S.W. helped create expression constructs and performed the infectivity assay using the MLV-based pseudoviruses. W.S. carried out the S1 shedding experiment and purified antibodies. H.Z. and M.G.K. performed the flow cytometry experiment. K.R., C.B. and, D.R.W. purified polyclonal IgG samples and contributed anti-S antibody expression constructs. All authors analyzed the data. B.C. and M.G.K. wrote the manuscript with input from all other authors.

## Competing Interests

All authors declare no competing interests.

## Data Availability

All data generated during and/or analyzed during the current study are available from the corresponding authors on reasonable request.

## Notes

### Competing Interest Statement

The authors have declared no competing interest.

