## Supplementary materials for "Antibody-mediated cell entry of SARS-CoV-2"

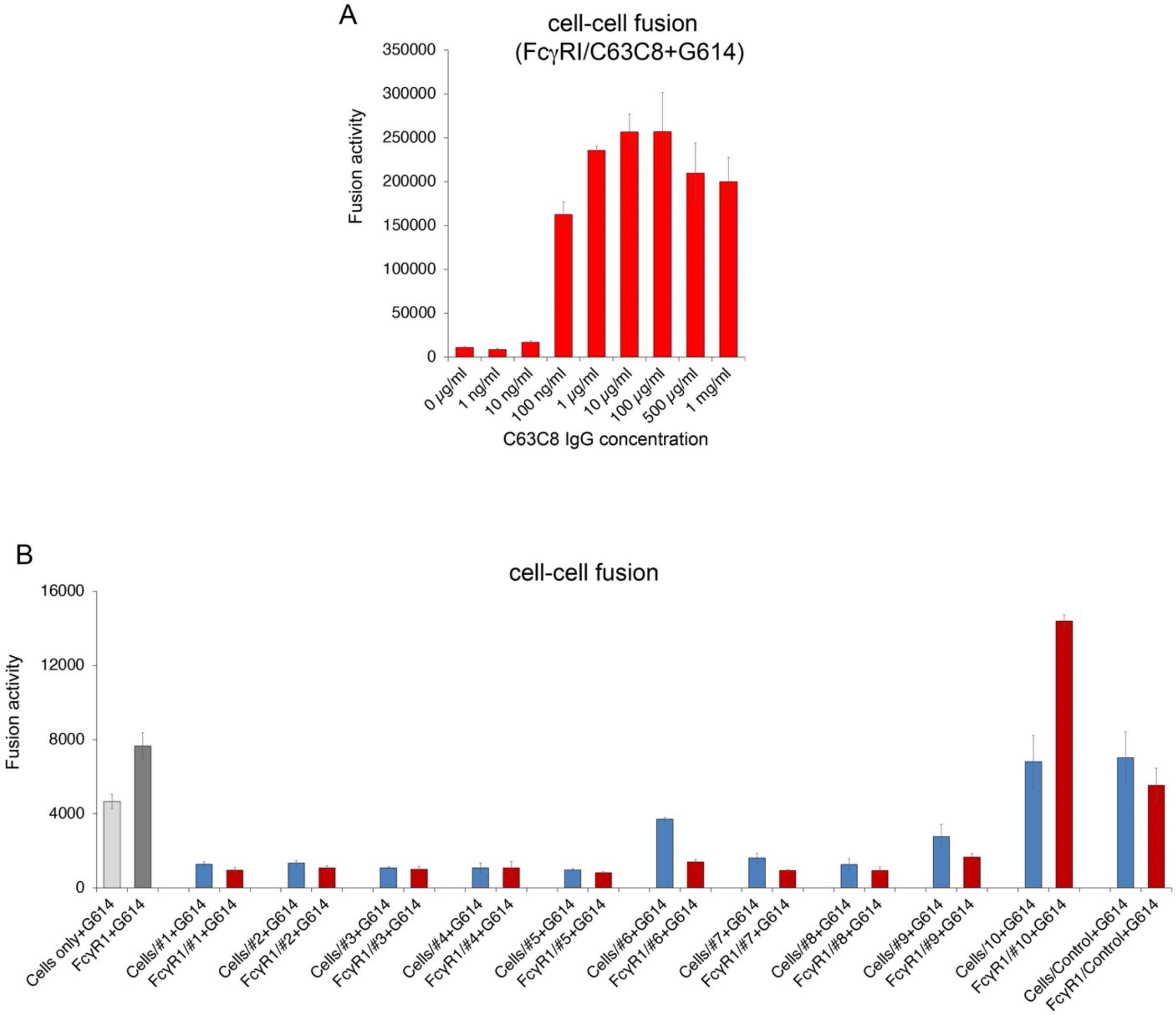


| Serum Sample | | SARS-CoV-2: G614 | |
|---|---|---|---|
| Sample # | Bleed Date | $ID_{50}$ | $ID_{80}$ |
| 1 | 27-May-2020 | 5,004 | 1,010 |
| 2 | 9-Jun-2020 | 4,529 | 653 |
| 3 | 16-Jun-2020 | 1,141 | 394 |
| 4 | 17-Jun-2020 | 4,261 | 753 |
| 5 | 16-Jul-2020 | 3,537 | 548 |
| 6 | 21-Jan-2021 | 1,538 | 721 |
| 7 | 19-Feb-2021 | 4,309 | 2,071 |
| 8 | 23-Feb-2021 | 764 | 210 |
| 9 | 5-Apr-2021 | 8,554 | 2,986 |
| 10 | 27-Apr-2021 | 7,463 | 2,202 |

**Figure S1. Membrane fusion mediated by antibody C63C8 and patient serum samples.** (**A**) Titration of C63C8 concentration in the assay for antibody/FcγRI mediated cell-cell fusion. HEK293T cells expressing FcγRI were decorated with C63C8 IgG at various concentrations and tested for membrane fusion with the full-length G614 S protein expressing cells. Cell-cell fusion led to reconstitution of α and ω fragments of β-galactosidase yielding an active enzyme and thus the fusion activity was quantified by a chemiluminescent assay. (**B**) The cell-cell fusion assay was used to analyze serum samples from ten convalescent individuals, collected between May 2020 and April 2021. A serum sample for an uninfected healthy individual was used as a control. The $ID_{50}$ and $ID_{80}$ values were determined using a pseudovirus-based neutralization assay[23].

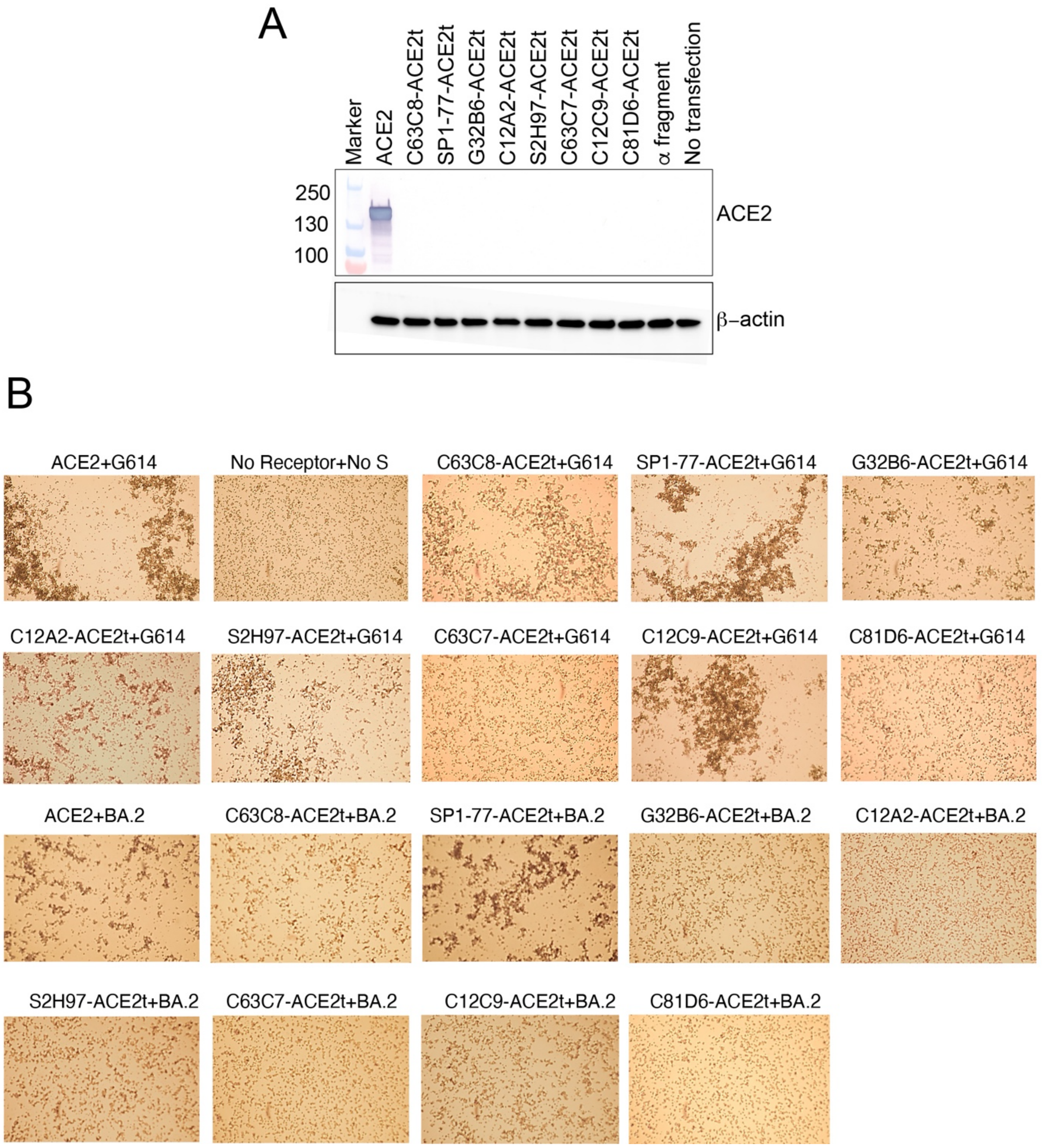


**Figure S2. Expression of endogenous ACE2 and cell association by mixing S-expressing and receptor-expressing cells.** (**A**) Expression of the endogenous ACE2 was monitored, by western blot using an antibody recognizing the catalytic domain of ACE2, when cells were transfected with various antibody-ACE2t constructs. Bands for the wildtype ACE2 and the sample loading control β-actin are indicated. (**B**) Images of cell association 15 min after S-expressing and receptor-expressing cells were mixed, as indicated.

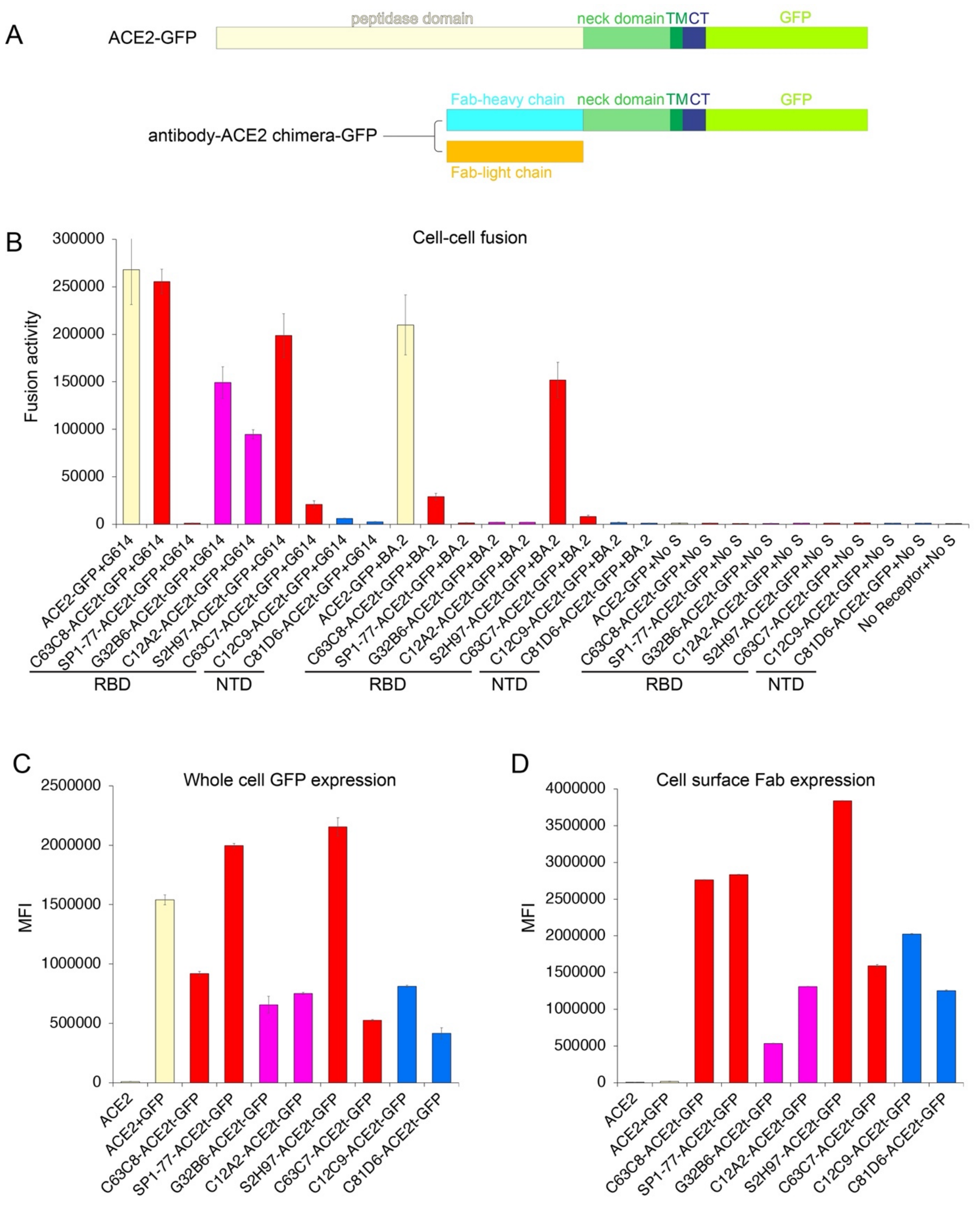


**Figure S3. Membrane fusion and expression of antibody-ACE2 chimeras.** (**A**) Schematic representation of the C-terminally GFP-tagged full-length human ACE2 and antibody-ACE2t expression constructs. (**B**) HEK293T cells transfected with GFP-tagged ACE2 and eight different

GFP-tagged antibody-ACE2t constructs were tested for membrane fusion with the full-length S protein (G614 or Omicron subvariant BA.2) expressing cells in the β-galactosidase-based cell-cell fusion assay. The GFP-tagged ACE2 was a positive control and no receptor/no S a negative control. (**C**) Expression levels of GFP-tagged constructs were quantified by flow cytometry. (**D**) Expression levels of GFP-tagged constructs were quantified using an anti-Fab secondary antibody conjugated with a dye by flow cytometry.

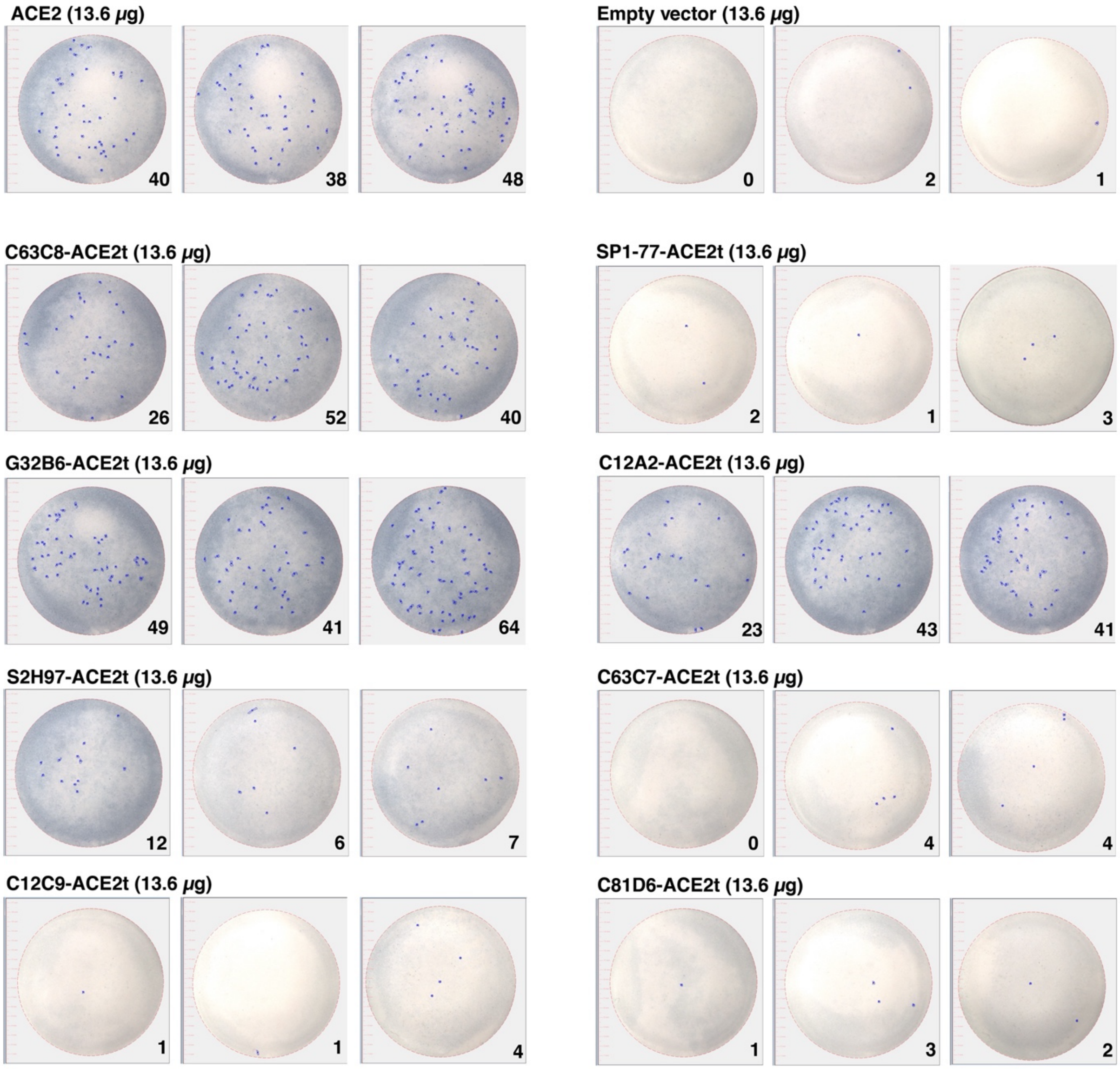


**Figure S4. Foci images of receptor-like antibody expressing MDCK cells infected by the authentic SARS-CoV-2.** Nonpermissive MDCK cells were first transfected with ACE2 or various antibody-ACE2t constructs and subsequently infected with the authentic SARS-CoV-2 G614 (B.1) virus at MOI of 0.005. Virus foci as a cluster of cells expressing viral antigen were imaged and counted using AID vSpot ELISPOT reader. The number of virus foci (dark spots) automatically identified by AID vSpot is shown in the lower corner of each well.

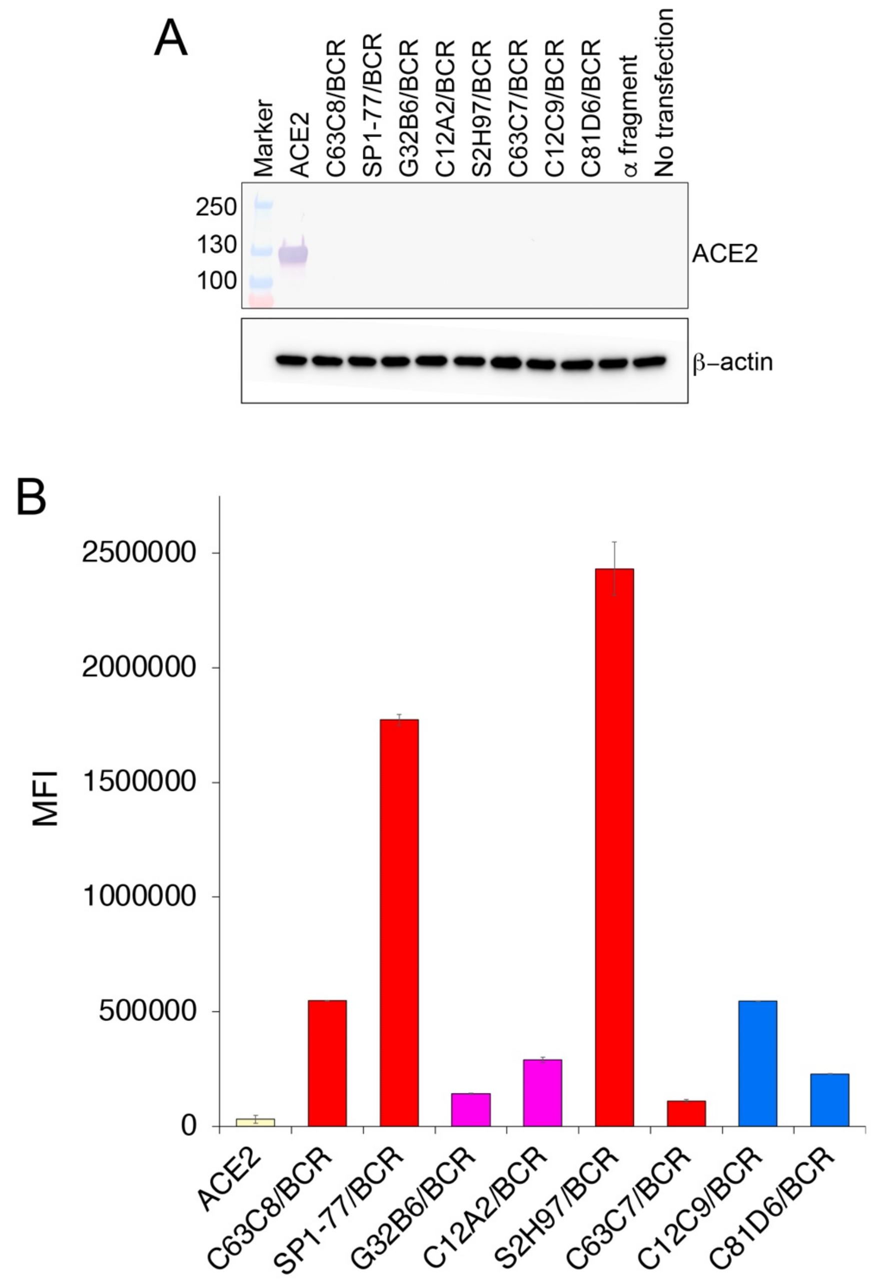


**Figure S5. Expression of endogenous ACE2 and various BCR constructs.** (**A**) Expression of the endogenous ACE2 was monitored, by western blot using an antibody recognizing the catalytic domain of ACE2, when cells were transfected with various BCR constructs. Bands for the wildtype ACE2 and the sample loading control β-actin are indicated. (**B**) Expression levels of BCR constructs were quantified using an anti-Fab secondary antibody conjugated with a dye by flow cytometry.

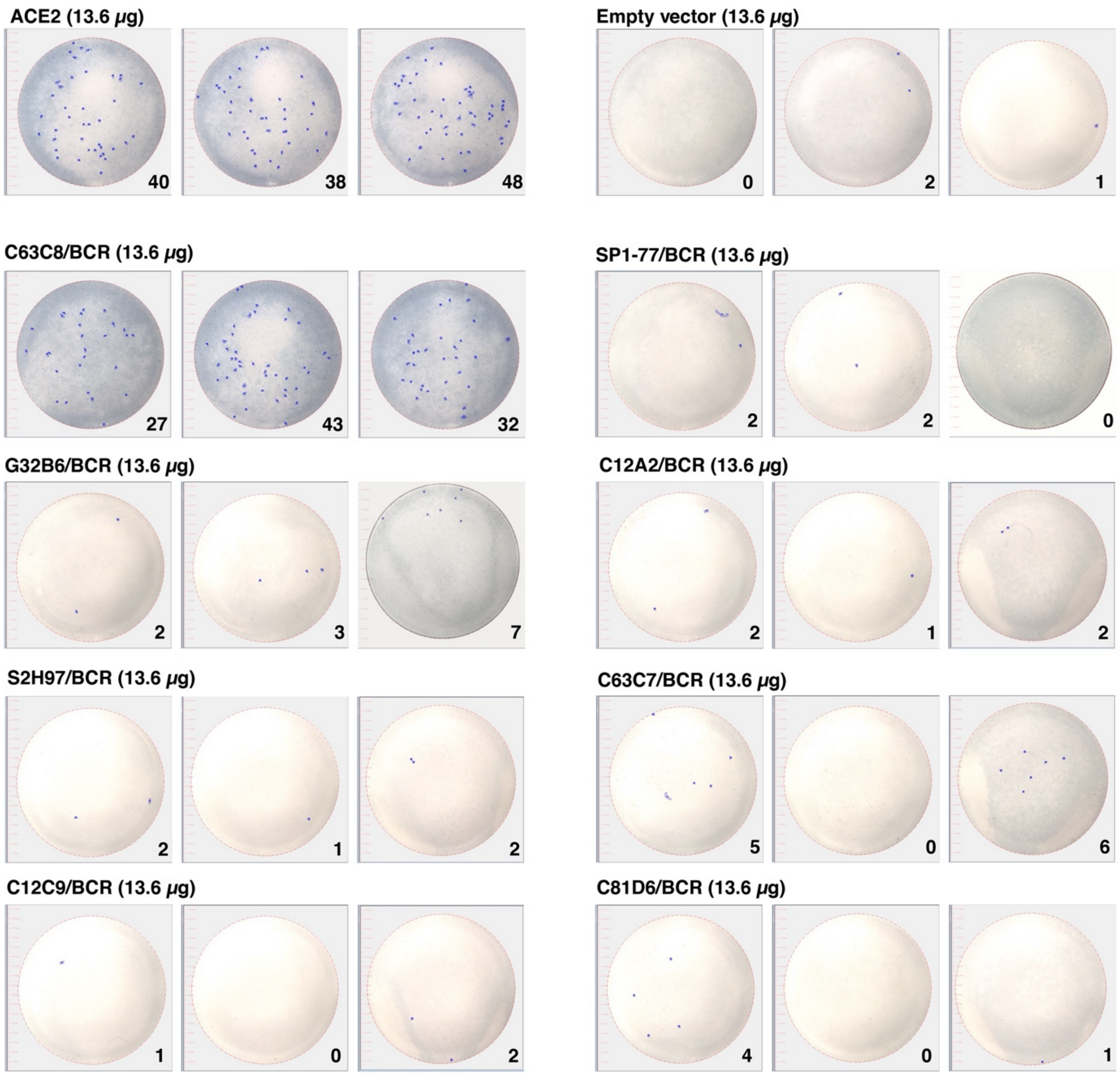


**Figure S6. Foci images of receptor-like antibody expressing MDCK cells infected by the authentic SARS-CoV-2.** Nonpermissive MDCK cells were first transfected with ACE2 or various BCR constructs and subsequently infected with the authentic New York-G614 (B.1) virus at MOI of 0.005. Postinfection supernatants were titrated by a focus-forming assay. Foci as a cluster of cells expressing viral antigen were imaged and counted using AID vSpot Spectrum. The number of foci (dark spots) automatically identified by AID vSpot is shown in the lower corner of each well.

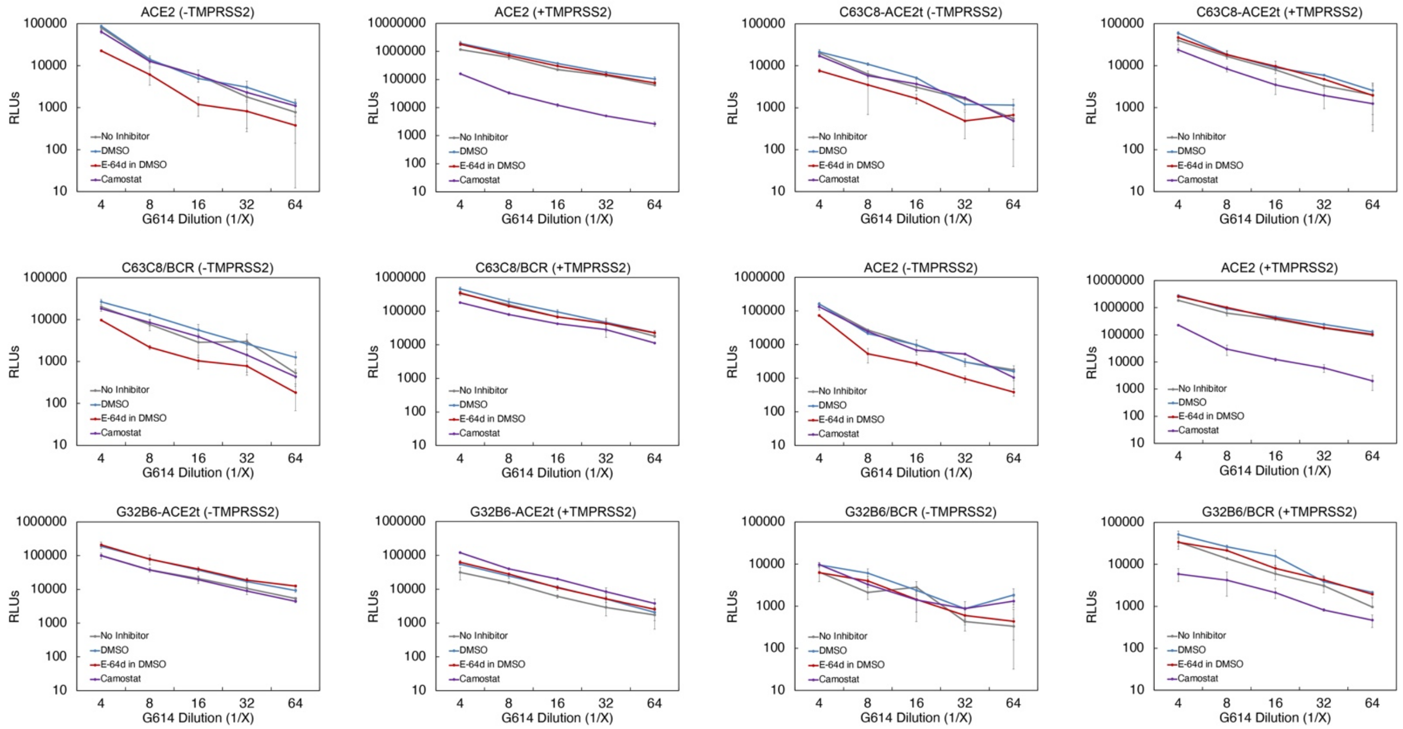


**Figure S7. Inhibition of SARS-CoV-2 entry mediated by receptor-like antibody.** Inhibition of viral infectivity by protease inhibitors. Pseudoviruses infection of HEK293T cells transfected with ACE2 or antibody constructs with/without TMPRSS2 was treated with E-64d (a cathepsin L inhibitor) or Camostat (TMPRSS2 inhibitor). DMSO, organic solvent used to dissolve E-64d.

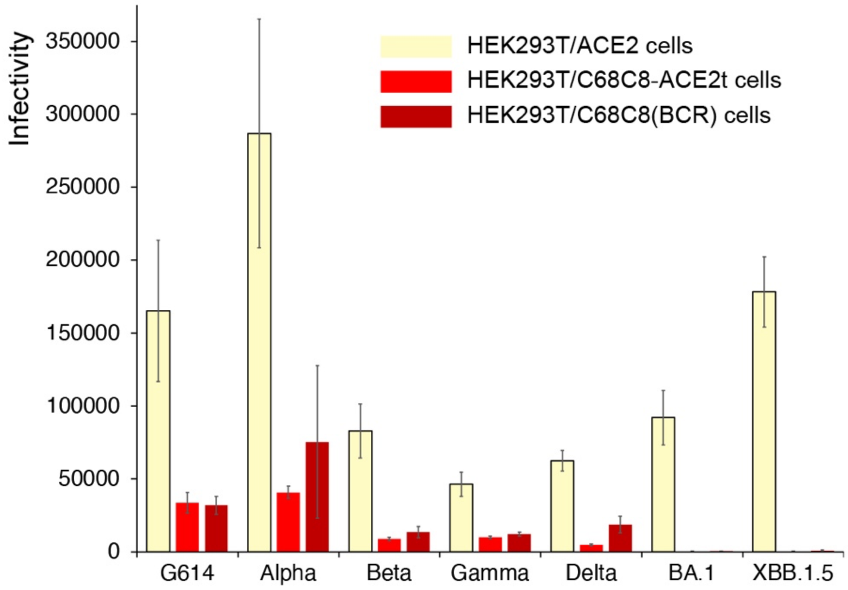


**Figure S8. Inhibition of SARS-CoV-2 entry mediated by receptor-like antibody.** C63C8-ACE2t and C63C8/BCR mediate infection by MLV-based pseudoviruses of various SARS-CoV-2 variants of concern.

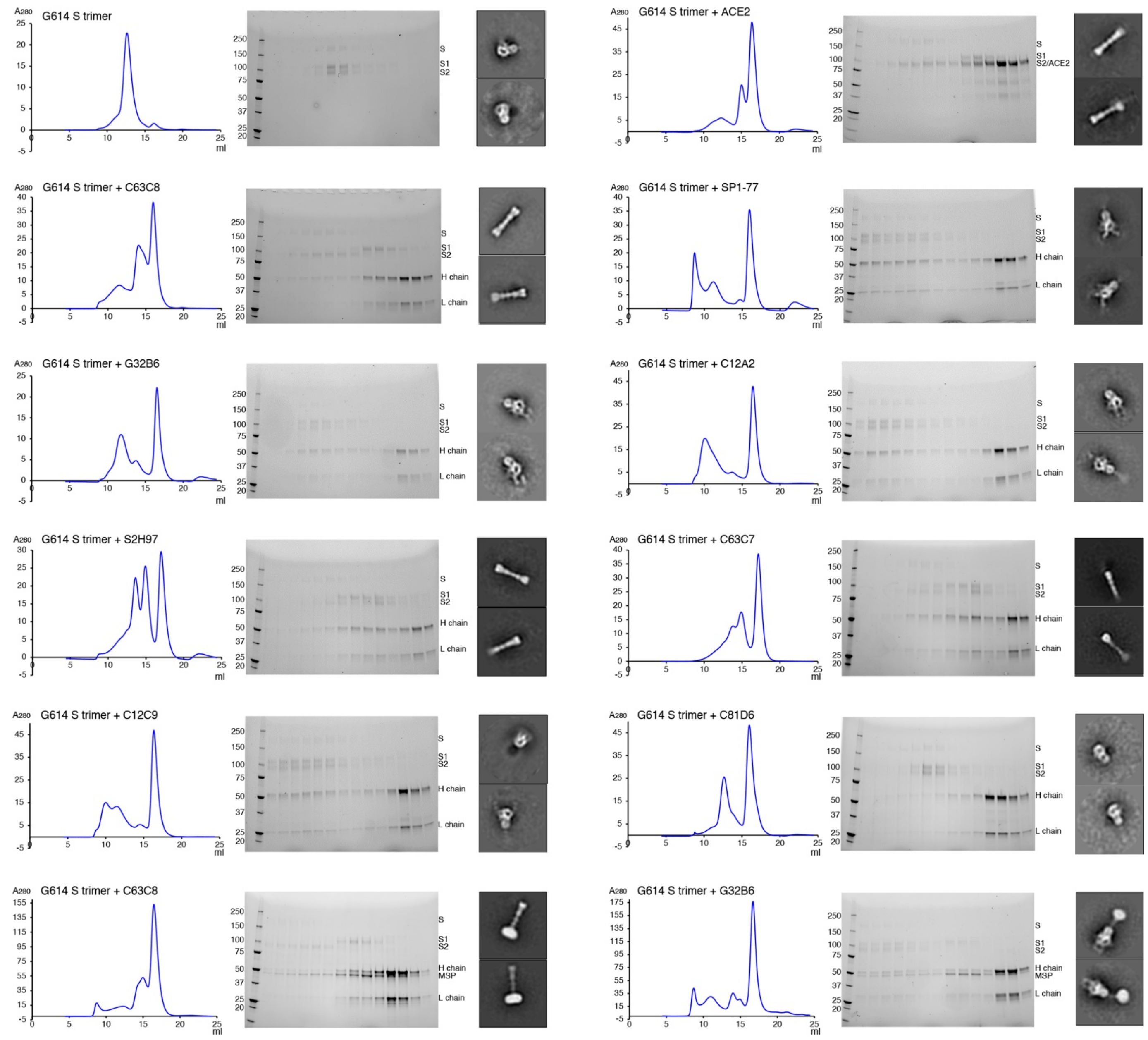


**Figure S9. S1 dissociation induced by ACE2 or monoclonal antibodies.** The purified full-length G614 S trimer solubilized in detergent was incubated with soluble ACE2 or various monoclonal IgG antibodies as indicated and resolved by gel-filtration chromatography on a Superose 6 column. Peaks fractions were analyzed by SDS-PAGE with bands for the uncleaved S, ACE2, S1 fragment and S2 fragment, as well as heavy (H) and light (L) chains for antibody indicated. Representative 2D averages by negative stain EM of the peak fractions are also shown. The box size of 2D averages is ~880Å. Bottom row, the G614 S trimer was reconstituted in lipid nanodiscs, incubated with either C63C8 or G32B6, and resolved by gel-filtration chromatography. Peaks fractions were analyzed by SDS-PAGE with bands for the uncleaved S, S1 fragment and S2 fragment, MSP (membrane scaffold protein) as well as heavy (H) and light (L) chains for antibody indicated. Representative 2D averages by negative stain EM of the peak fractions are also shown. The box size of 2D averages is ~880Å.

**Table S1. Characteristic overview of mAb-ACE2t chimeras and BCR**

| mAb | Spike Epitope | | Construct | Cell-cell association | | Cell-cell fusion | | HIV-based pseudovirus | | Authentic virus |
|---|---|---|---|---|---|---|---|---|---|---|
| | Tong[a] | Barnes[b] | | G614 S | BA.2 S | G614 S | BA.2 S | G614 S | BA.1 S | G614 |
| ACE2 | RBD-2 | | | +++ | +++ | +++ | +++ | +++ | +++ | +++ |
| C63C8 | RBD-1 | Class 3 | ACE2t | ++ | + | +++ | + | ++ | - | +++ |
| | | | BCR | ND | ND | +++ | - | +++ | - | +++ |
| SP1-77 | RBD-1 | Class 3 | ACE2t | +++ | +++ | - | - | - | - | - |
| | | | BCR | ND | ND | - | - | - | - | - |
| G32B6 | RBD-2 | Class 1/2 | ACE2t | ++ | - | +++ | - | +++ | - | +++ |
| | | | BCR | ND | ND | ++ | - | - | - | - |
| C12A2 | RBD-2 | Class 1/2 | ACE2t | ++ | - | +++ | - | ++ | - | +++ |
| | | | BCR | ND | ND | ++ | - | - | - | - |
| S2H97 | RBD-3 | Class 4 | ACE2t | ++ | + | +++ | +++ | + | - | + |
| | | | BCR | ND | ND | +++ | +++ | - | - | - |
| C63C7 | RBD-3 | Class 4 | ACE2t | - | - | + | + | - | - | - |
| | | | BCR | ND | ND | - | - | - | - | - |
| C12C9 | NTD-1 | NA | ACE2t | +++ | - | - | - | ++ | - | - |
| | | | BCR | ND | ND | - | - | - | - | - |
| C81D6 | NTD-2 | NA | ACE2t | - | - | - | - | - | - | - |
| | | | BCR | ND | ND | - | - | - | - | - |

[a]Nomenclature from Tong et al[45]; [b]Nomenclature from Barnes et al[40]. +++, Strong; ++, Medium; +, Weak; - No activity; ND, not determined; NA, not applicable.
